# A kinetically-driven free exchange mechanism of EmrE antiport sacrifices coupling efficiency in favor of promiscuity

**DOI:** 10.1101/141937

**Authors:** Anne E. Robinson, Nathan E. Thomas, Emma A. Morrison, Bryan Balthazor, Katherine A. Henzler-Wildman

## Abstract

EmrE is a small multidrug resistance transporter found in *E. coli* that confers resistance to toxic polyaromatic cations due to its proton-coupled antiport of these substrates. Here we show that EmrE breaks the rules generally deemed essential for coupled antiport. NMR spectra reveal that EmrE can simultaneously bind and cotransport proton and drug. The functional consequence of this finding is an exceptionally promiscuous transporter: Not only can EmrE export diverse drug substrates, it can couple antiport of a drug to either one or two protons, performing both electrogenic and electroneutral transport of a single substrate. We present a new kinetically-driven free exchange model for EmrE antiport that is consistent with these results and recapitulates ΔpH-driven concentrative drug uptake. Our results suggest that EmrE sacrifices coupling efficiency for initial transport speed and multidrug specificity.

**SIGNIFICANCE:** EmrE facilitates *E. coli* multidrug resistance by coupling drug efflux to proton import. This antiport mechanism has been thought to occur via a pure exchange model which achieves coupled antiport by restricting when the single binding pocket can alternate access between opposite sides of the membrane. We test this model using NMR titrations and transport assays and find it cannot account for EmrE antiport activity. We propose a new kinetically-driven free exchange model of antiport with fewer restrictions that better accounts for the highly promiscuous nature of EmrE drug efflux. This model expands our understanding of coupled antiport and has implications for transporter design and drug development.

## INTRODUCTION

Secondary active transport moves one substrate across a membrane against its concentration gradient by coupling it to downhill transport of a second substrate, often a proton. This coupled transport process may move both substrates in the same direction (symport) or in opposite directions (antiport). To move molecules across the membrane, the substrate binding site must be alternately accessible to either side of the membrane. Symport or antiport of two substrates is generally explained using models that restrict this alternating access of the transporter to specific states (Fig. 1). These models are appealing because they provide a simple mechanism that efficiently couples transport of the two substrates.

**Fig. 1.**
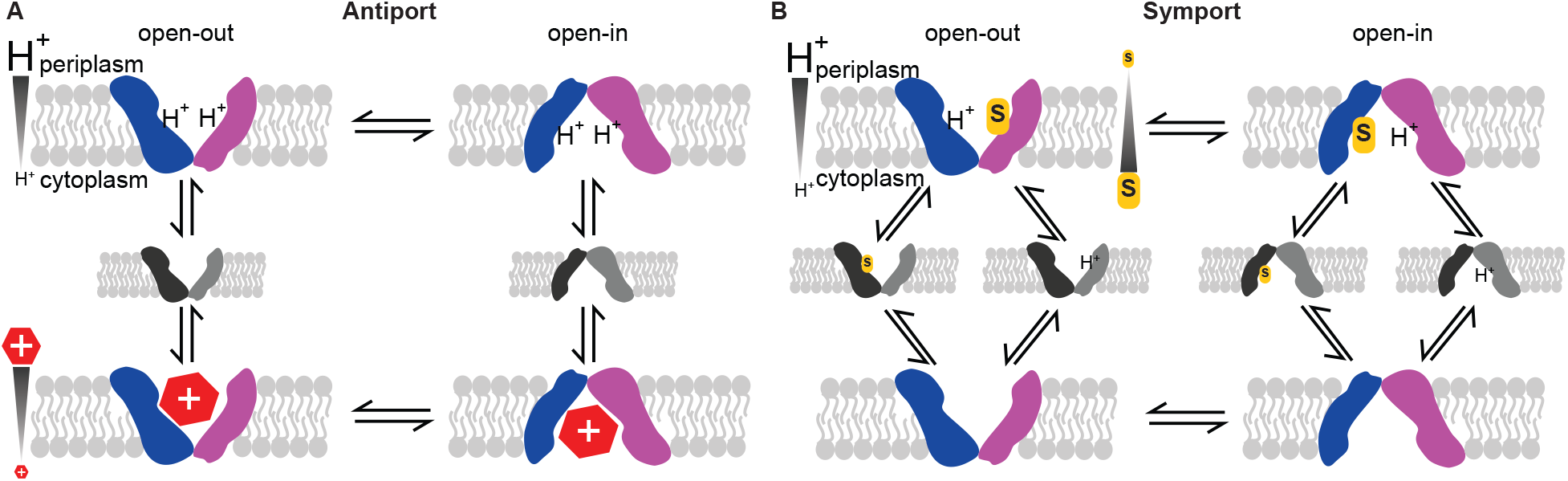
Models of tightly-coupled secondary transport. A) The pure exchange model of coupled antiport applied to EmrE (red hexagon, TPP^+^). The asymmetric homodimer of EmrE is represented by the distinct shapes of the two conformations, and the two monomers are colored blue and magenta. B) General model for proton-coupled symport of a substrate (yellow S) illustrating the differences in which states participate in alternating access. Grey structures represent intermediates which are restricted from alternating access in these models.

Here we investigate the mechanism of proton/drug antiport by the small multidrug resistance transporter, EmrE. EmrE uses the proton motive force (PMF) across the inner membrane of *E. coli* to drive efflux of toxic polyaromatic cations, conferring resistance to these compounds. The single binding pocket of EmrE is defined by two glutamate 14 residues (1–3), one on each of the two monomers in the asymmetric homodimer (4–7). This binding site can accommodate one drug-substrate or up to two protons. Alternating access of the asymmetric homodimer is achieved by a conformation swap between the two monomers (5) and is necessary for transport activity (8). Traditionally, EmrE antiport has been explained by the pure exchange model (9) in figure 1A. Such “pure exchange” of one drug for two protons with no slippage results in tightly coupled stoichiometric antiport. This is achieved by 1) limiting substrate binding such that both substrates never bind simultaneously and 2) limiting in/out exchange (alternating access) to substrate bound states (fully protonated or drug-bound). Several lines of evidence support this model. Competition between drug and proton binding is demonstrated by substrate-induced proton release (10, 11), a decrease in substrate-binding affinity at low pH (1) and a bell-shaped pH-dependence of transport activity (3). In addition, observation of electrogenic transport of monovalent, but not divalent, substrates is consistent with a 2:1 H^+^/drug transport stoichiometry (12). Other transport mechanisms have been considered previously (11), but the traditional pure exchange model has been favored in the absence of compelling data to justify selection of a more complex scheme.

However, the highly dynamic nature of EmrE (13–15), critical for its ability to transport diverse substrates, is hard to reconcile with the strict limitations on alternating access in the pure exchange model. Recent NMR data has provided evidence that EmrE violates at least the second stipulation of the traditional model: protonation of drug-free EmrE is asymmetric (16) such that a singly protonated state exists near neutral pH, and all of the protonation states (2H^+^-bound, 1H^+^-bound, empty) engage in alternating access (16, 17). These findings suggest the need to develop a new model for EmrE transport activity. In this study, we use NMR spectroscopy and liposomal flux assays to test the pure-exchange model of EmrE antiport. We show that EmrE violates both requirements of pure-exchange antiport and utilizes multiple drug:proton antiport stoichiometries. We develop a new kinetically-driven “free exchange” EmrE antiport model which reconciles the novel states and unrestricted alternating access behavior of EmrE with its well-established proton-driven drug efflux activity. Our model suggests that the coupling efficiency of H^+^/drug antiport is sacrificed for the ability of EmrE to efficiently efflux diverse substrates.

## RESULTS

### Substrate binding is not exclusive

Competition between drug and proton binding to EmrE is well-established, but the data do not prove mutually exclusive binding, a stipulation of the pure exchange model (1, 10, 11). The recent demonstration of asymmetry in proton binding (16) led us to reconsider whether EmrE can bind a drug and proton simultaneously. To test this we performed NMR pH titrations of EmrE saturated with the drug substrate tetraphenylphosphonium (TPP^+^) and solubilized in isotropic bicelles. We have previously shown that E14 and H110 are the only residues in EmrE that titrate near neutral pH (16). If proton and drug binding are exclusive, as predicted by the traditional model of EmrE antiport, then TPP^+^-saturated EmrE should not bind any protons and thus will not titrate with pH. However, many peaks in the NMR spectrum do titrate with pH (Fig. 2A and S1), demonstrating that protonation does occur when TPP^+^ is bound. In contrast to the two protonation events observed for drug-free EmrE (pKa values 7.0 ± 0.1 and 8.2 ± 0.3) (16), the linear movement of the peaks in the TPP^+^-bound NMR pH titration is consistent with a single protonation event with a pKa of 6.8 ± 0.1 (Fig. 2B and S2). To test whether this protonation occurs on the critical E14 residue, we repeated the pH titration with TPP^+^-saturated E14D-EmrE, which has a lower pKa (1–3, 16). As expected, we observed a shift in the titration midpoint to lower pH, reflecting the lower pKa of E14D-EmrE (Fig. 2A and S3) and confirming we are monitoring protonation of E14 in drug-bound EmrE.

**Fig. 2.**
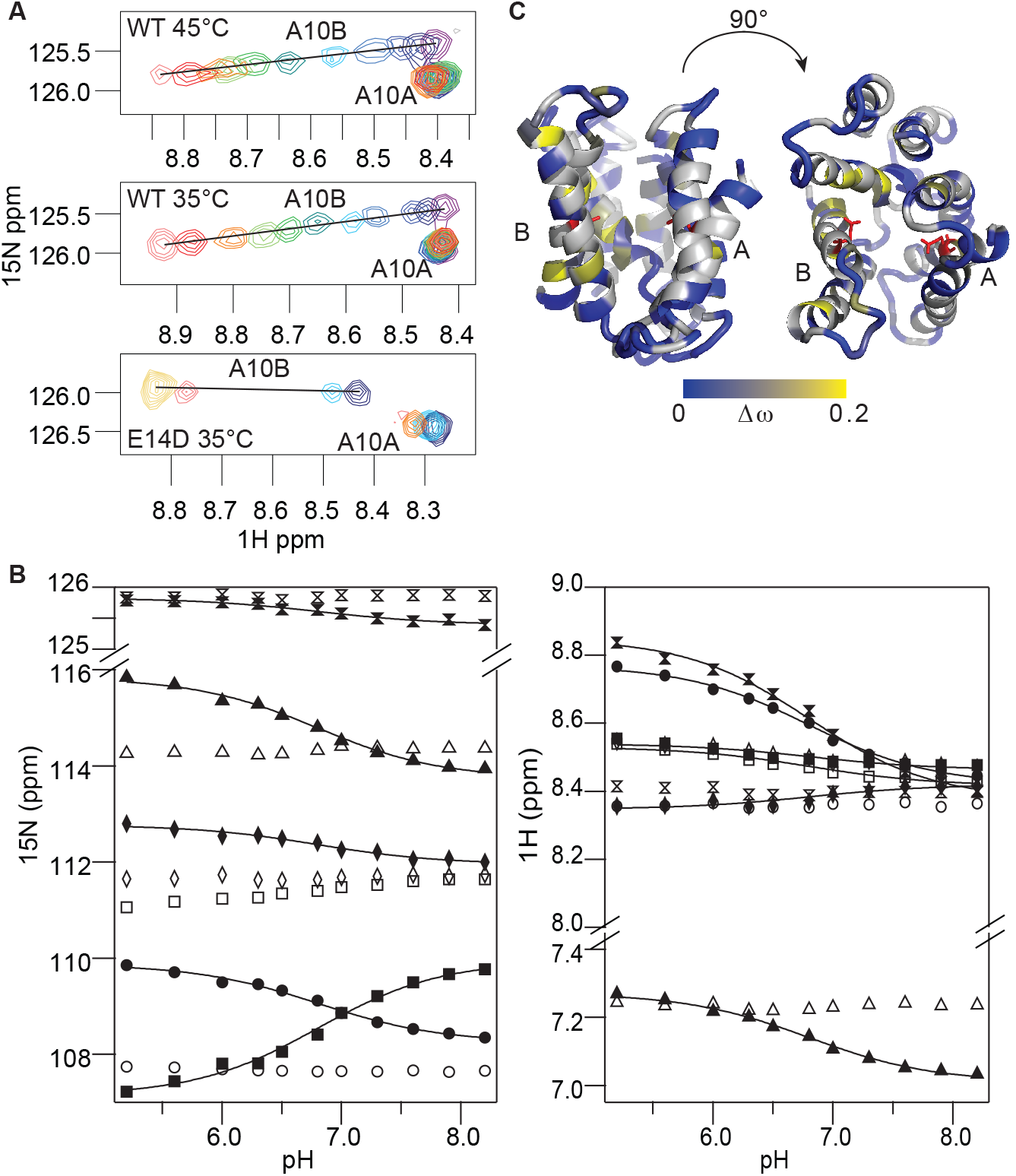
EmrE can bind drug and proton simultaneously. A) NMR pH titrations of TPP^+^-bound ^2^H/^15^N-WT (45 °C and 35 °C) and E14D (35 °C) EmrE shown for residue A10 (Full spectra in Fig. S1 and S3). pH values range from 5.2 (pink) to 8.3 (purple) for WT and 4.0 (yellow) to 8.0 (navy) for E14D-EmrE. B) Global fitting of the pH-dependent chemical shifts from monomer B residues (solid symbols) yields a single pKa of 6.8 ± 0.1 at 45 °C. Monomer A residues (open symbols) have relatively pH-independent chemical shifts. Error bars are smaller than the symbols. C) Plotting the chemical shift changes (Δω) between pH 5.6 and 7.6 onto the structure of TPP^+^-bound EmrE (PDB 3B5D) highlights the localization of pH-dependent effects in monomer B (grey, residues not resolved at both pH values). E14 is shown as red sticks.

It is well known that TPP^+^ and H^+^ binding is competitive such that the TPP^+^ binding affinity of EmrE is weaker at low pH (1). Thus, it is important to ensure that the pH-dependent chemical shift changes are not due to pH-dependent loss of substrate binding. We used ITC to measure TPP^+^ binding at low pH. In isotropic bicelles at 45 °C (matching the NMR sample conditions), the *K*_D_^apparent^ for TPP^+^ is 70 ± 9 μM at pH 5.5 (Table S1). With this affinity, >95% of EmrE will remain TPP^+^-bound in the NMR sample at pH 5.5. As a second more direct test, we performed an NMR-monitored TPP^+^ titration at pH 5.2. As expected, there were no chemical shift changes with increasing TPP^+^ concentrations (Fig. S4). Therefore, the pH-induced NMR spectral changes are not due to loss of TPP^+^ binding. The simplest explanation for the NMR data is that EmrE can simultaneously bind a drug and a proton at physiological pH.

### Substrate binding is asymmetric

Mapping the residues that sense E14 protonation onto the structure reveals a broad distribution (Fig. 2C), consistent with protonation-dependent conformational changes in drug-bound EmrE. This is not surprising given the protonation-dependent transport activity of EmrE (3) and the coupled structural and dynamic changes that occur upon protonation in the absence of substrate (16–18). Interestingly, residues corresponding to monomer B in the asymmetric homodimer have much larger chemical shift changes upon pH titration of TPP^+^-bound EmrE (Fig. 2C). This implies that TPP^+^ binds asymmetrically to the E14 on monomer A, while the E14 on monomer B remains accessible for protonation. This is further supported by the TPP^+^ titration at low pH which shows that residues corresponding to monomer A are more sensitive to the concentration of TPP^+^ (Fig. S4). Such asymmetric substrate binding was suggested by the cryoEM structure of TPP^+^-bound EmrE (6). It is also consistent with the asymmetric structure of the EmrE homodimer as demonstrated by the unique chemical shifts of the two E14 residues (19) and the asymmetric protonation of the two E14 residues in drug-free EmrE (16).

### Drug binding only releases one proton at low pH

To validate this extraordinary finding that an antiporter can simultaneously bind both substrates, we measured TPP^+^-induced proton release from EmrE at low pH. According to the pure exchange model, substrate binding is mutually exclusive such that both protons should be released from the EmrE homodimer upon TPP^+^ binding at low pH. In contrast, if EmrE is able to bind TPP^+^ and a proton simultaneously, TPP^+^ binding will only trigger release of a single proton per dimer at low pH. A previous measurement of TPP^+^-induced proton release was inconclusive because the amount of TPP^+^ added was insufficient to saturate EmrE at low pH (10). We repeated this measurement with EmrE solubilized in isotropic bicelles using a saturating TPP^+^ concentration and observed release of 1.2 ± 0.2 protons per EmrE dimer at pH 5.5 (Fig. S5). Because the weak buffering required for direct proton detection in this assay results in relatively large errors, we used a second experimental approach to verify the results. By measuring TPP^+^ binding with ITC in multiple buffers with different heats of ionization (20) we could determine the number of protons released per TPP^+^ binding event and confirm binding saturation (Table S1). We again detected 1.2 ± 0.1 protons released per dimer and confirmed a 1:1 TPP^+^/dimer binding stoichiometry. These proton release values are much closer to 1 than 2, consistent with the NMR data demonstrating simultaneous binding of 1 TPP^+^ and 1 H^+^ to the EmrE dimer at low pH.

### Alternating access of EmrE bound to both substrates

The ability of EmrE to bind a drug and proton simultaneously will only affect net transport if this state engages in alternating access. For EmrE, the two monomers within the asymmetric homodimer swap conformations to switch between open-in and open-out (alternate access), and this can be quantitatively measured using TROSY-selected ZZ exchange NMR experiments (5, 21). We compared the rate of alternating access for TPP^+^-saturated EmrE in isotropic bicelles at high pH (only drug bound) and low pH (both drug and proton bound) and found they were nearly identical. (Fig. 3, S6). Thus, EmrE can move both substrates across the membrane at the same time, violating the expected behavior of an antiporter, and requiring the development of a new transport model.

**Fig. 3.**
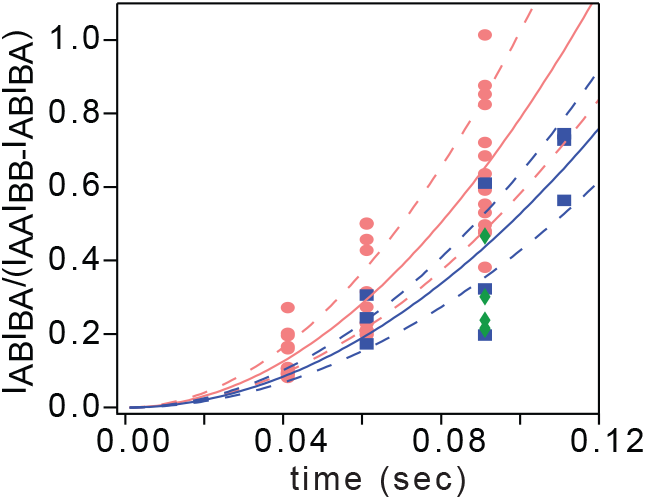
All EmrE substrate-bound states engage in alternating access. The composite peak ratio from ^1^H-^15^N TROSY-selected ZZ-exchange NMR experiments is shown for multiple residues at pH 5.2 (pink circles), pH 7.0 (green diamonds), and pH 8.0 (blue squares). Global fits yield alternating access rates of 8.9 ± 1.2 s^−1^ at pH 5.2 (TPP^+^/H^+^ both bound) and 7.3 ± 0.7 s^−1^ at pH 8.0 (only TPP^+^ bound). Error was estimated with jackknife analysis of individual residue fits (dotted lines).

Importantly, the peaks in the NMR spectra correspond to the two distinct monomer *conformations* (shapes in Fig. 1A), not monomer identities (colors in Fig. 1A). Thus, TPP^+^ preferentially binds to the monomer in *conformation* A. Since the two monomers swap conformations during alternating access, this means that TPP^+^ (and H^+^) is (are) swapped back and forth between the two monomers as part of the alternating access process. Therefore, although only one E14 is necessary for TPP^+^ binding, the swapping of substrate between monomers during alternating access will require both E14 residues for transport. This is consistent with both the “functional symmetry” of E14 (22) and the dominant negative phenotype of E14 mutants *in vivo* (23).

### A new kinetically-driven free exchange model for EmrE transport

To accommodate the new states and transitions of EmrE, we expanded the transport scheme (Fig. 4A), removing the restrictions on simultaneous substrate binding and alternating access that are imposed by the pure exchange model. This model is *kinetically-driven:* net flux will be determined by the relative rates of individual steps. To understand if it is possible to produce the well-established proton/drug antiport activity of EmrE with this scheme we performed mathematical simulations. We can estimate all of the rate constants from our own and others’ experimental data (Fig. S7 and Table S2). Rates of alternating access are based on NMR dynamics measurements (5, 16, 17). TPP^+^ on- and off-rate estimates were determined previously for detergent-solubilized EmrE (11). We assumed fast proton on-rates (≈10^10^ M^−1^s^−1^) and used the pKa values determined by NMR (this work and (16)) to estimate off-rates. These are reasonable assumptions given the small size of H^+^, the relatively fast on-rates for TPP^+^ binding, and the fact that binding affinities of drug-substrates are primarily determined by the off-rate (11).

Numerical simulation of ΔpH-driven transport (Appendix 1) results in rapid concentrative uptake of TPP^+^ into liposomes (Figure 4B). A 100-fold pH gradient drives 80-fold concentration of TPP^+^, a modest reduction in coupling efficiency compared to the pure exchange model with a strict 2H^+^:1TPP^+^ transport stoichiometry. These results clearly demonstrate that ΔpH-driven coupled antiport of TPP^+^ can be achieved in our new kinetically-driven model without restricting alternating access or substrate binding.

**Fig. 4.**
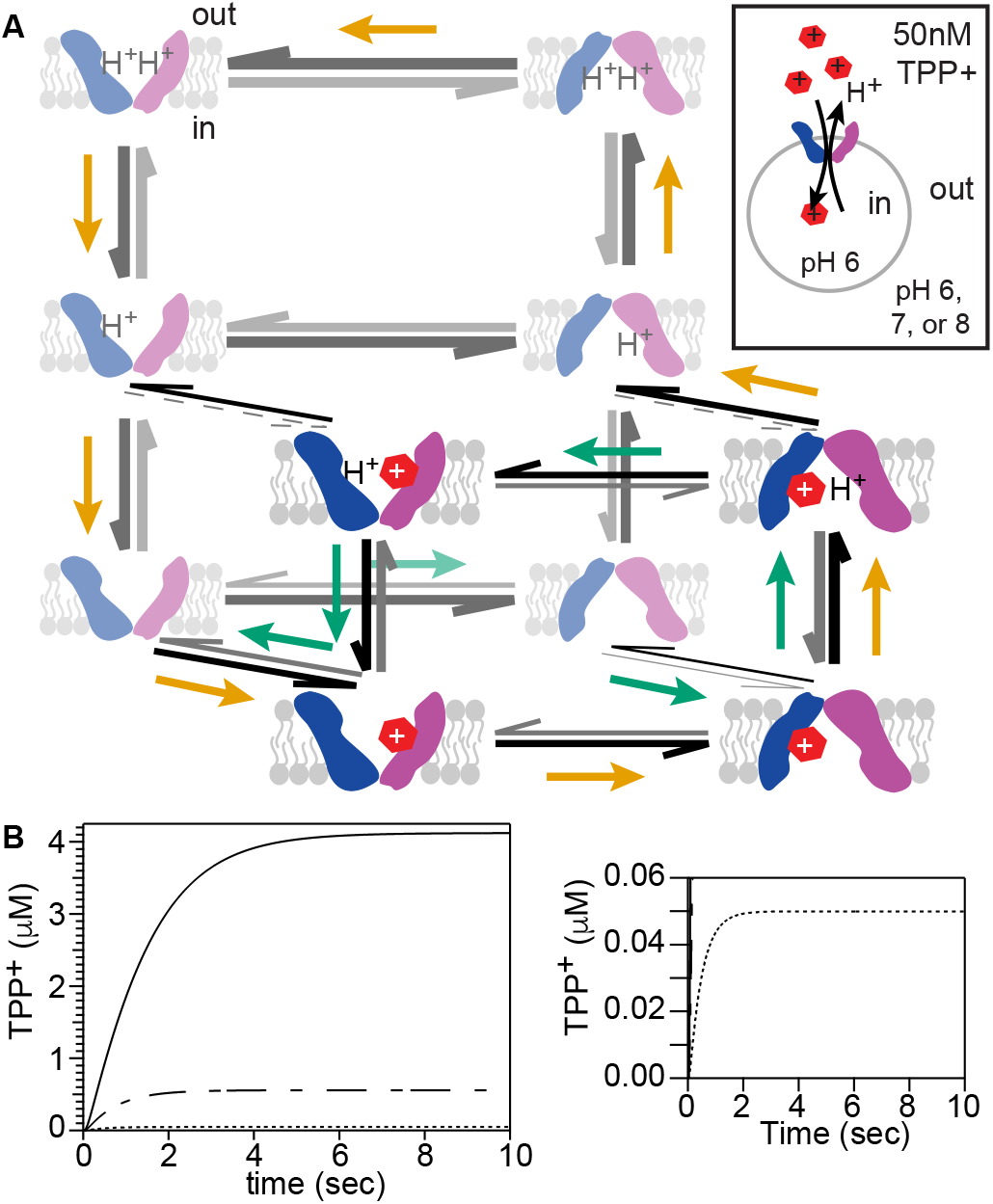
Simulation of the kinetically-driven free exchange model. A) Transport scheme including all known states of EmrE. Inset illustrates the liposomal loading that was modeled. Line weights reflect the relative frequency of each transition during the Gillespie simulation (pH 8 out, pH 6 in); darker lines indicate higher frequency. Orange arrows highlight the most probable antiport path; blue arrows highlight the symport path observed when the model is run with hypothetical positive linkage. Dotted lines, transition did not occur during the simulation time. B) Kinetic simulation of TPP^+^ uptake into liposomes using the rate constants in Fig. S7 and Table S2 results in proton-driven concentrative uptake of TPP^+^ into EmrE proteoliposomes (pH 6 inside) when ΔpH=1 (pH 7 outside, dashed lines) or ΔpH=2 (pH 8 outside, solid lines). When ΔpH=0 (dotted line, expanded to right), 50 nM TPP^+^ rapidly equilibrates.

To better understand how coupled antiport occurs in our new model, we developed a single molecule Gillespie simulation (Appendix 2) (24). Unlike the previous deterministic simulation, this stochastic simulation ignores the finite liposomal volume, but informs on the relative frequency of each transition, which is indicated by arrow thickness in Fig. 4A. Interestingly, TPP^+^ is most likely to bind proton-free EmrE but be released subsequent to protonation. Simultaneous proton and TPP^+^ binding enhances the release of TPP^+^ (Fig. S7), resulting in faster turnover. This mechanism may be advantageous for a multidrug transporter because it allows efficient release of substrates with a wide range of affinities.

While our simulations demonstrate coupled antiport, future work will be needed to make them quantitatively accurate. Several rate constants were estimated using simplifying assumptions, and all were measured using solubilized EmrE. Importantly, solubilization results in a symmetric environment rather than the asymmetric conditions experienced in a membrane under a proton motive force. Nevertheless, the ability of our simulation to qualitatively recapitulate EmrE antiport is noteworthy because it breaks long-standing assumptions about the mechanism of coupled antiport.

Unlike the pure exchange model, our model allows *free exchange* of the transporter, with alternating access permitted in all states. This has significant functional implications because it allows for *free exchange* of substrates with multiple transport stoichiometries. This differs dramatically from the single antiport stoichiometry achieved with pure exchange of substrates across the membrane. To test our kinetically-driven free exchange model, we turned to an experimental transport assay.

### EmrE can perform both 2:1 and 1:1 H^+^/drug antiport

To test if EmrE can indeed perform antiport with multiple transport stoichiometries, we performed liposomal transport assays. EmrE has previously been shown to perform 2:1 H^+^/drug^+^ transport (12) and we repeated that experiment, confirming 2:1 transport in our EmrE proteolipomes (data not shown). Here we chose to monitor the countertransported proton, not the drug substrate because the key question is how many protons are transported per drug substrate. We reconstituted EmrE into proteoliposomes with a strongly-buffered pH 6 interior and weakly-buffered pH 8 exterior, creating a pH gradient to drive drug uptake (Fig. 5A). We monitored release of protons upon addition of a monovalent polyaromatic cation substrate to the external solution. A 2:1 H^+^/drug^+^ transport stoichiometry is electrogenic and will result in rapid charge buildup, preventing significant transport. In contrast, a 1:1 H^+^/drug^+^ transport stoichiometry is electroneutral and should proceed under all conditions (Fig. 5A). To distinguish between these two transport stoichiometries, we included KCl both inside and outside the liposomes and tested whether valinomycin, an ionophore that allows K^+^ flux across the membrane to dissipate any charge buildup, was necessary for transport.

**Fig. 5.**
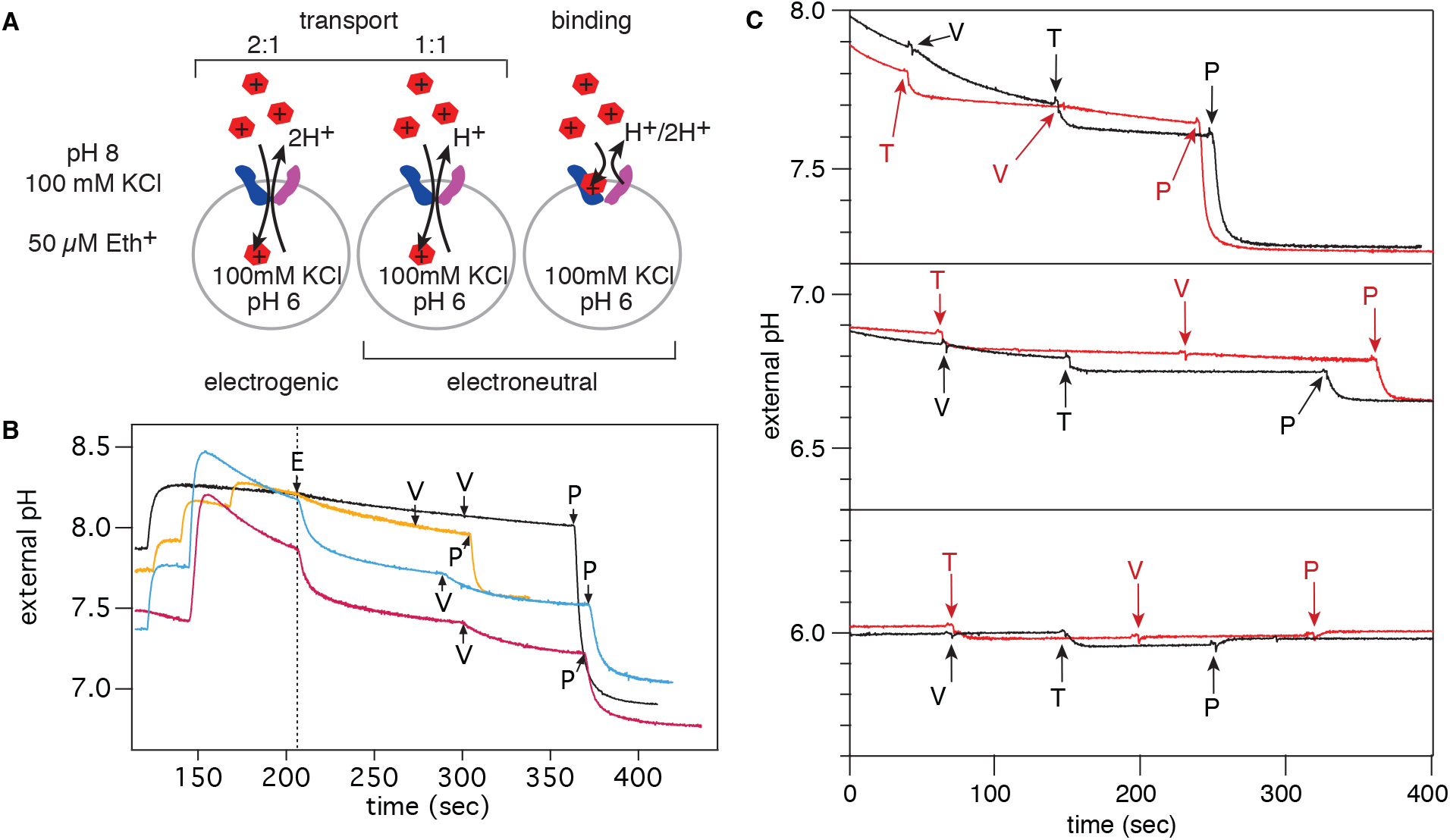
EmrE transports Eth^+^ and TPP^+^ with multiple H^+^/drug stoichiometries. A) ΔpH-driven drug uptake by EmrE proteoliposomes was monitored by direct measurement of the countertransported proton. Cartoon showing the Eth^+^ uptake assay conditions illustrates the potential sources of proton release detected in the assay. B) In the presence of a pH gradient, addition of 50 μM Eth^+^ (E) outside results in a sudden release of protons prior to addition of valinomycin (V), indicating an electroneutral process. Proton release is EmrE-dependent as demonstrated by DCCD inactivation of EmrE (yellow) and empty liposomes (black). The same proton release is observed for 1:500 (red) and 1:1000 (blue) EmrE:lipid mole ratios, showing that proton release is due to electroneutral transport and not binding. Addition of a protonophore (P) confirms the proton gradient was maintained. Proton release was quantified using the observed pH shift upon addition of a known aliquot of NaOH. C) The same assay using 40 μM TPP^+^ (T) as the drug substrate. Black and red traces differentiate T and V order of addition. Proton release from the proteoliposomes always occurs upon addition of T, not V, indicating an electroneutral process. Repeating the experiment with external pH of pH 6, 7, or 8 (bottom to top) shows the effect of different ΔpH (0, 1, 2, respectively) on the T-induced proton release. More protons are released at higher external pH (greater ΔpH) confirming that 1:1 transport is occurring. Representative traces (see Tables S3 and S4 and Fig. S8 for additional replicates, controls, and quantitation).

Ethidium (Eth^+^), an EmrE substrate with a weaker binding affinity *(K_D_* at least 1000-fold weaker (25)) and decreased hydrophobicity compared to TPP^+^, is more well-suited to transport assays. Upon addition of Eth^+^ to EmrE proteoliposomes, substantial proton release is observed in the absence of valinomycin (Fig. 5B). The Eth^+^-induced proton release is EmrE-dependent as it is blocked by EmrE inhibition with DCCD and is not observed for empty liposomes (Fig. 5B). Addition of protonophore at the end of each assay confirms the integrity of the proteoliposome pH gradient (Fig. 5B). Thus, the valinomycin-independent proton release is consistent with an electroneutral process.

Both proton efflux due to 1:1 H^+^/Eth^+^ antiport and proton release upon Eth^+^ binding are electroneutral processes (Fig. 5A). To confirm the signal was not simply due to Eth^+^ binding, we repeated the transport assay with double the protein:lipid ratio but the same total lipid. This results in twice as much protein, so proton release should double if it is due to Eth^+^ binding. However, since the size and number of liposomes is unchanged, the same steady-state transport equilibrium will be reached and the measured proton release should not change if it is due to 1:1 transport. The quantity of protons released is the same (Fig. 5B and Table S3), demonstrating that the majority of the proton release signal is due to 1:1 H^+^/Eth^+^ antiport.

If EmrE utilizes both 1:1 and 2:1 transport stoichiometries, then addition of the ionophore valinomycin should allow 2:1 transport and drive further proton release. In fact, additional proton release was observed upon subsequent addition of valinomycin (Fig. 5B). Thus, Eth^+^ is transported with both a 1:1 and a 2:1 H^+^/drug transport stoichiometry.

We repeated these transport assays with the high affinity substrate, TPP^+^, used for the NMR experiments. As with Eth^+^, we observed that TPP^+^ transport occurs regardless of whether valinomycin is present (Fig. 5C). As before, non-specific proton leakage through the liposome is excluded because a steady state is reached without full dissipation of the proton gradient (Figure 5C), as shown by a further pH drop upon addition of FCCP. The TPP^+^-induced proton release is EmrE-dependent as it is blocked by EmrE inhibition with DCCD (Figure S8).

To distinguish between proton release upon TPP^+^ binding and electroneutral antiport we determined the effect of external pH on the number of protons released. Increasing external pH increases ΔpH, and thus the driving force for 1:1 antiport. Therefore, proton efflux coupled to TPP^+^ uptake (1:1 antiport) will be *greater* at higher pH_external_. However, proton release upon drug binding will *decrease* at higher pH_external_ (1) since fewer protons are bound to EmrE initially. More protons are liberated at higher external pH (greater ΔpH): 46 ± 2 nmol H^+^/ml at pH 7.7_ext_, ΔpH=1.7 versus 31 ± 2 nmol H^+^/ml at pH 7.0_ext_, ΔpH=1 (Table S4). Unlike with Eth^+^, subsequent addition of valinomycin does not result in additional proton release (Fig. 5C). TPP^+^ is a high affinity ligand (25). 1:1 transport likely yields a TPP^+^_internal_ concentration that is sufficient to saturate open-in EmrE such that no additional net flux can occur.

To confirm that the signal is not solely a result of TPP^+^ binding, we repeated the assay with twice the protein:lipid ratio. As before, this doubles the protein concentration while keeping liposome internal and external volume constant. Proton-release due to binding will increase in proportion to the EmrE concentration while transport should be unchanged. We observe an increase of 1.2 nmol H^+^ released per dimer, from 2.3 nmol H^+^ released per dimer to 3.5 nmol H^+^ released per dimer (Table S4). The increase must be due to binding, demonstrating that there is measurable binding-induced proton release (1.2 nmol H^+^ per dimer) when using the tight-binding TPP^+^ for transport assays. However, it does not account for the total number of protons liberated from the liposomes upon TPP^+^ addition, and the remainder of the protons must come from 1:1 transport. These results show that TPP^+^ can also be transported with a 1:1 H^+^/drug stoichiometry, supporting our new kinetically-driven free exchange model for EmrE transport activity.

## DISCUSSION

### Reassessing the mechanism of secondary active transport

Advances in *in vitro* studies of transporter function and structure have expanded our understanding of the range of conformational changes that can produce alternating access (26, 27), and the mechanisms controlling stoichiometry and transport (28). Recently, two transporters, MdfA and PepTST, have been found to use different proton/substrate transport stoichiometries when transporting substrates of different size or charge (29, 30). Here we show that EmrE can transport a *single* substrate with multiple proton/substrate transport stoichiometries. This flexibility of transport stoichiometry is achieved by a number of unexpected features. First, TPP^+^ binds asymmetrically in the active site, making space for a proton to bind simultaneously. Second, all of the EmrE proton- and/or drug-bound states are capable of engaging in alternating access. These phenomena of mutual binding and unlimited alternating access at first seem to jeopardize EmrE’s capacity for coupled antiport, but a closer look reveals the efficacy and benefits of such a mechanism.

How is antiport driven with these unique parameters? In this kinetically-driven model, the relative rates of the individual steps will determine the efficiency of proton-coupled transport. Although H^+^ and TPP^+^ can bind simultaneously, their binding is still negatively linked, resulting in differential TPP^+^ affinity for the open-in and open-out states of EmrE in the presence of a transmembrane pH gradient. In fact, a pH gradient may even skew the equilibrium between the open-in and open-out states (17). Such differential substrate affinity is important for determining the relative kinetics and efficiency of the transport cycle, particularly in the presence of leak pathways (31). Our new free exchange model relies on both TPP^+^/H^+^ binding competition and the thermodynamic and kinetic asymmetry introduced by the PMF to drive productive transport by EmrE. This has an interesting parallel to the finding that the PMF controls the rate of chemiosmotically-driven LacY symport (32). Many of the features of EmrE are also reminiscent of the *de novo* designed transporter, Rocker (33), although EmrE is more efficiently coupled. It suggests that functionally coupled transport can be achieved without the need to invoke significant constraints on the states and transitions of the transporter, perhaps providing new insights for the rational design of *de novo* transporters.

### Biological implications of the new mechanism

The free exchange model has an elegant simplicity of its own, placing no structural or dynamic constraints upon EmrE. This is functionally relevant because the highly dynamic nature of EmrE is important for its promiscuous multidrug recognition (13–15). Our model suggests that simultaneous drug and proton binding may even be advantageous, speeding up the release of tight binding substrates that would otherwise compromise EmrE’s ability to rapidly pump toxic molecules out of *E. coli*. The free exchange model allows for extreme adaptability in a minimalistic protein, sacrificing coupling efficiency for increased transport speed (Fig. S7). Interestingly, reduced coupling efficiency was previously proposed as a necessary compromise for multisubstrate specificity (34, 35), and reduced efficiency may be an acceptable tradeoff in the context of a bacterial cell that continuously regenerates the PMF, similar to futile ATP hydrolysis by P-glycoprotein (36, 37).

Pure exchange models only allow pure exchange of the counter-transported substrates and are appealing due to their apparent simplicity, tight coupling and stoichiometric antiport. However, even a simple shared-carrier model, which allows for alternating access of the apo transporter, will give rise to coupled antiport (9). The data presented here takes this one step further, demonstrating that the promiscuous transporter, EmrE, can move both substrates across the membrane simultaneously. This has been assumed to be behavior indicative of a symporter, and not possible for an antiporter. The flexibility inherent in our free exchange model accommodates the observed ease of converting SMR transporters from antiporters to symporters (38–40) and suggests that the kinetically-driven mechanism may be common across the SMR family. Since this kinetically-driven model relies upon the relative rates of substrate binding and alternating access, and the transported substrate determines the rate of alternating access (25), it may be possible to design substrates which would convert an SMR pump from an antiporter to a symporter or vice versa.

In fact, a single transporter acting as both a symporter and antiporter of different substrates has been reported for W63G-EmrE (38). W63G-EmrE performs proton-coupled antiport of erythromycin but symport of bis-tris-propane *in vitro* and *in vivo*, conferring resistance to erythromycin but performing concentrative uptake of bis-tris-propane into *E. coli* to toxic levels (38). This unusual phenotype cannot be explained by the classical models of proton-coupled transport, which place mutually exclusive requirements on alternating access of the drug-free transporter (Fig. 1). However, the behavior of W63G-EmrE is readily explained with our kinetically-driven free exchange model. Negative linkage between proton and erythromycin binding will favor independent binding of the two substrates and antiport via a shared-carrier model. On the other hand, positive linkage between bis-tris-propane and proton binding will favor simultaneous drug and proton binding and symport. In our kinetic simulations, switching the negative linkage between drug and proton binding observed for WT EmrE to positive linkage by altering both the proton and drug on- and off-rates can switch WT EmrE from an antiporter to a (relatively inefficient) symporter (Fig. 4A). Future experiments will be needed to more thoroughly test whether robust symport can be achieved. This will likely vary with each SMR because the interplay between the relative rates of all the microscopic steps will be important for the efficiency and efficacy of either antiport or symport.

In an era when antibiotic resistance and drug-delivery pose a serious challenge, our results suggest a novel strategy. If an MDR efflux pump can indeed function as both a symporter and antiporter, and this balance can be shifted by properties of the transported substrate, perhaps it can be subverted to drive drugs *in* to bacteria, providing a new route for drug delivery. Despite decades of study, EmrE continues to reveal its surprising complexity and expand our understanding of basic transport mechanisms and multidrug efflux.

## METHODS

### EmrE expression and purification

EmrE was expressed in *E. coli* BL21(DE3) and purified as described previously (5) using a pET15b vector with an N-terminal 6x His tag kindly provided by G. Chang. For ^2^H/^15^N-labelled EmrE, the M9 media contained 1 g ^15^NH_4_Cl, 2 g glucose, 0.5 g ^15^N,D Isogro (Sigma, St. Louis, MO), and 1 multivitamin per liter D_2_O. Purification was via Ni-NTA chromatography followed by size exclusion chromatography with a Superdex 200 column equilibrated in either NMR buffer (for bicelle NMR samples) or the appropriate inside buffer (for liposomal reconstitution and transport assays), each with 10 mM decyl maltoside (DM, Anatrace, Maumee, OH).

### NMR sample preparation and data acquisition

EmrE in NMR buffer (20 mM acetate, 100 mM MOPS, 100 mM bicine) with 10 mM DM was reconstituted into DMPC (1,2-dimyristoyl-sn-glycero-3-phosphocholine, Avanti Polar Lipids, Alabaster, AL) at 75:1 lipid:EmrE monomer mole ratio following the protocol in (41). EmrE proteoliposomes were collected by ultracentrifugation (100,000 g, 2 hr, 6 °C) and resuspended in NMR buffer with DHPC (1,2-dihexanoyl-sn-glycero-3-phosphocholine, Avanti Polar Lipids, Alabaster, AL) and freeze-thawed 3 times to create q=0.33 (47) bicelles. Final NMR samples contained 0.5–1.0 mM EmrE monomer, 10% D_2_O, 0.05% NaN_3_, 2 mM TCEP (*tris*(2-carboxyethyl)phosphine), 2 mM EDTA (Ethylenediaminetetraacetic acid), and 2 mM DSS (4,4-dimethyl-4-silapentane-1-sulfonic acid). EmrE concentration was determined using the extinction coefficient reported previously (38370 l mol^−1^ cm^−1^) (5).

The pH of NMR samples was measured at the experimental temperature in a water bath with a pH electrode calibrated at the same temperature and adjusted by step-wise addition of weak HCl or NaOH. When possible, pH titrations were performed with two samples at high and low pH that were mixed to titrate pH.

All NMR spectra were collected on a 700 MHz Varian Inova spectrometer equipped with a room temperature probe. ^1^H chemical shifts were referenced with DSS, ^15^N chemical shifts were referenced indirectly, and temperature was calibrated using ethylene glycol. pH titrations were collected with ^1^H-^15^N BEST-TROSY-HSQC pulse sequences (44, 45). The TROSY-selected ZZ exchange experiment (21) was modified and run as previously described (5). All data were processed with NMRPipe (42) and analyzed in CcpNmr analysis (43). pKa values were determined by fitting proton and nitrogen chemical shifts as a function of pH to the following equation (48):

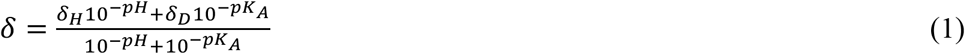

ZZ-exchange rates were determined by globally fitting a composite ratio of the auto and cross peak intensities for all residues with well-resolved cross- and auto-peaks in the ZZ-exchange spectra, with error determined by jackknife analysis of individual residue fits, as described previously (5, 46):

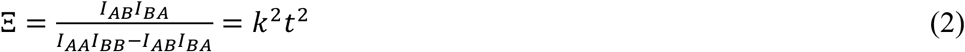

### ITC experiments

EmrE was reconstituted into DLPC/DHPC isotropic bicelles with effective q-values of 0.33 and a minimum ratio of 100:1 DLPC:EmrE. Additional lipid was added to low-concentration samples to keep the total lipid concentration above 40 mM and preserve bicellar morphology (47). EmrE is fully dimeric at all protein:lipid ratios used(49).

Titrations of TPP^+^ into EmrE were carried out in multiple buffers (Table S1) with a range of ionization enthalpies. Both TPP^+^ and EmrE solutions contained matching concentrations of isotropic bicelles, 20 mM buffer and 20 mM NaCl. 5 mM TPP^+^ was titrated into 835 μM EmrE in a TA Instruments Low Volume Nano calorimeter using the ITCRun software (TA Instruments, Lindon, UT) with 2.5 μL injections, stirring at 350 rpm at 45 °C. Sample pH was checked at 45 °C before and after each experiment. Data is reported in Table S1.

Buffer ionization enthalpies (50) were adjusted to 45 °C using the reported standard molar heat capacity change at 25 °C. Each titration was analyzed independently, confirming the 1 TPP^+^:EmrE dimer binding stoichiometry under all conditions. Plots of 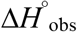 (determined by ITC) vs 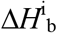 (ΔH of buffer ionization) have a slope of *nH*^+^, regardless of how complex a mechanistic model is considered, as shown below (eqns. 4,7,10).

Assuming that only two protons or one drug molecule can bind at a time (model in Fig. 1A), the simplifying condition of independent sites (no linkage) can be made. Following the notation of (20), for two independent and identical proton-binding sites, the system is represented by:

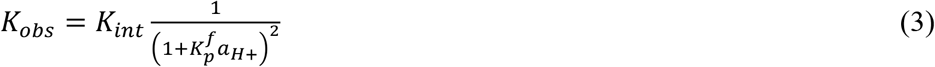

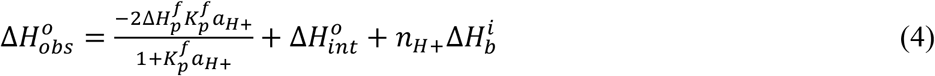

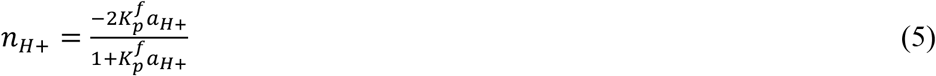

where *K*_obs_ is the observed TPP^+^ binding affinity and 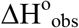 is the observed enthalpy change upon TPP^+^ binding as measured experimentally with ITC. *n*_H+_ is the change in the number of protons bound by EmrE upon TPP^+^ binding, *a*_H+_ is the proton activity (10^−pH^), *Kint* is the TPP^+^-association constant for apo EmrE, 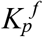 is the H^+^ binding constant for apo EmrE (10^pKa^), 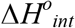 is the enthalpy change for TPP^+^ binding to apo EmrE, 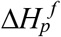 is the enthalpy of protonation of apo EmrE and 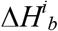 is the ionization enthalpy of the buffer.

For two independent and non-identical proton-binding sites, as reported for EmrE (16), the system is represented by:

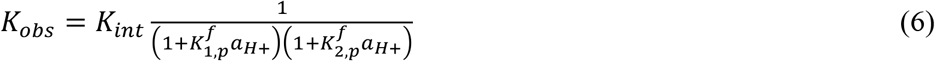

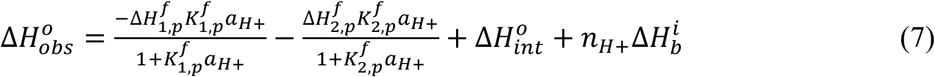

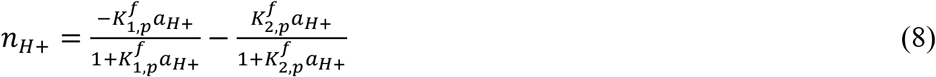

where 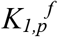 and 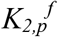 are the H^+^ binding constants for binding the first and second protons by apo EmrE (10^pKa1^, 10^pKa2^) and 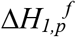 and 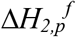 similarly represent the enthalpy of the first and second protonation steps for apo EmrE.

If, however, one proton and one drug molecule can bind simultaneously, the mechanism becomes more complex (Figure 6A). In the case that EmrE can bind two protons at independent non-identical sites, one drug molecule, or a single proton and a drug molecule, the system is represented by:

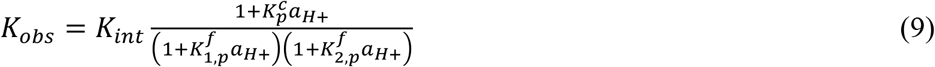

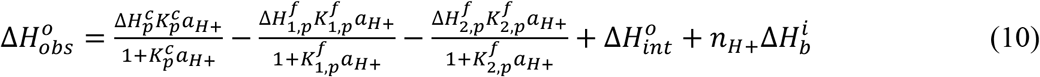

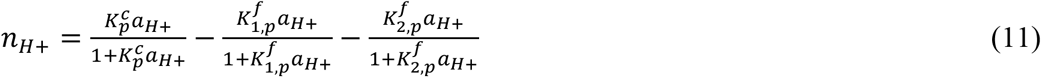

where 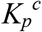 is the H^+^-binding constant for TPP^+^-bound EmrE and 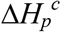 is the enthalpy of protonation for TPP^+^-bound EmrE. The NMR data presented in figure 2 do not show any indication that EmrE can bind TPP^+^ and 2 H^+^ simultaneously, so we did not consider more complex scenarios.

### Proton release measurements

EmrE was prepared in q=0.33 DLPC:DHPC isotropic bicelles as described above, but the final steps of isotropic bicelle preparation were performed with unbuffered solutions (20 mM NaCl, pH 7) to create a final sample with weak buffering capacity. Assays were performed with 3, 5, and 10 nmol EmrE and 80 mM total lipid in triplicate. The pH was monitored in real-time using a Biotrode micro pH electrode as a saturating concentration of TPP^+^ (0.9 mM) was added to the sample as determined from ITC measurements of the binding affinity. 10 nmol of NaOH was added at the end of the assay for quantitation. pH was recorded before TPP^+^ addition, after TPP^+^ addition and after HCl addition in order to fit a linear baseline to each segment to account for pH drift occurring in the weakly buffered solution. The number of protons released was calculated from the slope of a plot of the protons released versus the EmrE concentration. Error was propagated through each of the linear fitting steps to estimate the error in the number of protons released.

### Proteoliposome reconstitution for transport assays

POPE, POPC, or POPG polar lipids (Avanti Polar Lipids, Alabaster, AL) in chloroform were dried under nitrogen and rinsed twice with pentane or lyophilized overnight to remove residual chloroform. Dry lipids were hydrated at 20 mg/ml for 1 hr in the appropriate inside buffer for the desired assay (see below), extruded 21 times through a 0.2μm filter (Avanti Polar Lipids, Alabaster, AL) and permeabilized with 0.5% octyl-glucoside for 15 min at room temperature. Purified EmrE in 10 mM DM was added to obtain a 1:64 or 1:50 w/w protein:lipid mole ratio for ethidium or TPP^+^ transport assays, respectively, resulting in ≈1:1000 or ≈1:800 EmrE:lipid mole ratio. A second set of samples was reconstituted with twice the protein:lipid ratio for control experiments, and empty liposomes were prepared by adding 10 mM DM without protein. After 20 min incubation, detergent was removed with amberlite (Sigma, St. Louis, MO) as in NMR sample preparation. 8 ml of proteoliposomes were then dialyzed (Slide-A-Lyzer Dialysis Cassettes, 2K MWCO, Thermo Fisher Scientific, Waltham, MA) against 1 L inside buffer for 24 hours twice to remove any residual detergent. Final liposomes were aliquoted, flash frozen in liquid nitrogen and stored at −80°C until use. Empty liposomes were prepared in the same way, with 10 mM DM added to simulate the reconstitution process.

### Proton transport assay

EmrE was reconstituted into 3:1 POPE/POPG liposomes (TPP^+^ transport) or 3:1 POPC:POPG liposomes (ethidium^+^ transport) in strongly buffered high-potassium inside buffer (20 mM potassium phosphate, 300 mM KCl, pH 6 for TPP^+^ transport, or 100 mM MES, 100 mM KCl, pH 6 for ethidium^+^ transport) as described and extruded 21x through a 0.4 μm filter (TPP^+^ transport) or 0.2 μm filter (ethidium^+^ transport). Proteoliposomes were passed over two 10 mL Sephadex G25 desalting columns equilibrated with weakly buffered outside buffer (1 mM potassium phosphate, 300 mM KCl, pH 6 for TPP^+^ transport, or 0.5 mM MOPS, 300 mM KCl, pH 6 for ethidium^+^ transport). 1.2–1.4 ml total volume was used directly for each TPP^+^ transport assay. 1 mL of proteoliposomes was diluted with 0.5 mL outside buffer for each ethidium^+^ transport assay.

The external pH of the proteoliposome solution was recorded with a microelectrode using a Jenco analog pH meter, digitized with a DataQ data logger and recorded in real-time. External pH was adjusted with small aliquots of NaOH to the desired external starting pH and then substrate was added from a stock solution (38.5 mM TPP^+^ or 25 mM ethidium^+^) at the same external pH. The potassium ionophore valinomycin was added to 1 μg/ml to eliminate any charge build up due to electrogenic transport. The protonophore (carbonyl cyanide m-chlorophenyl hydrazine (CCCP) for Eth^+^ transport and carbonyl cyanide-p-trifluoromethoxyphenylhydrazone (FCCP) for TPP^+^ transport) was added to 1μg/ml to allow full pH equilibration at the end of the assay. A subset of EmrE proteoliposomes was inhibited by incubation with 0.5 mM N,N’-dicyclohexylcarbodiimide (DCCD) (300 mM stock in ethanol) at room temperature for 2 hr prior to use in the transport assay as a negative control. EmrE concentration was checked by A280 measurement using bicelles created by adding 4 mg DHPC to 1 ml proteoliposome stock followed by 3 freeze-thaw cycles.

### Building a Kinetic Simulation

We developed a system of non-linear differential equations to simulate EmrE transport activity based upon the schematic model presented in figure 4A. All rates were determined from experimental data presented in this and previous studies, as denoted in Table S2, except for the rate of proton binding and release. We assumed a diffusion-limited on-rate (10^10^ M^−1^s^−1^) for all proton-binding steps and used this assumption and our measured pKa values to calculate proton off-rates. This is a significant assumption, since the highly shifted pKa values demonstrate that E14 is not located in a typical environment exposed to bulk water. The model was simulated in Berkeley Madonna (version 9.0, Kagi shareware, Berkeley, CA) to model the amount of TPP^+^ loaded into the liposome over time under different conditions using the code in Appendix 1.

To appreciate the relative frequencies of each transition, the system of equations was modified to run a Gillespie simulation (24) using Octave (51) (Appendix 2). Here, total TPP^+^ accumulation was not counted or factored in. Rather, infinite inside and outside volumes were assumed (TPP^+^ concentrations were locked) and the frequency of each transition was recorded. Rates and terminology are defined in Table S2.

## ACKNOWLEDGEMENTS

We thank Geoff Chang for the EmrE expression plasmid. We thank Eric Galburt and John Robinson for assistance in developing the Gillespie transport model. This work was supported by the National Institutes of Health (1R01GM095839), NSF graduate research fellowships to E.M. and A.R. (DGE-1143954), and a Mr. and Mrs. Spencer T. Olin Fellowship for Women in Graduate Study to A.R.

